# Rice NIN-LIKE PROTEIN 3 plays a significant role in nitrogen use efficiency and grain yield under nitrate-sufficient conditions

**DOI:** 10.1101/2021.02.19.432039

**Authors:** Zi-Sheng Zhang, Jin-Qiu Xia, Alamin Alfatih, Ying Song, Yi-Jie Huang, Liang-Qi Sun, Guang-Yu Wan, Shi-Mei Wang, Yu-Ping Wang, Bin-Hua Hu, Guo-Hua Zhang, Peng Qin, Shi-Gui Li, Lin-Hui Yu, Jie Wu, Cheng-Bin Xiang

**Affiliations:** School of Life Sciences and Division of Molecular & Cell Biophysics, Hefei National Science Center for Physical Sciences at the Microscale, University of Science and Technology of China, The Innovation Academy of Seed Design, Chinese Academy of Sciences, Hefei, Anhui Province 230027, China; Rice Research Institute, Anhui Academy of Agricultural Sciences, Hefei 230031, China; Rice Research Institute, State Key Laboratory of Hybrid Rice, Sichuan Agricultural University, Chengdu, Sichuan, China; Biology Department, Brookhaven National Laboratory, Upton, New York, NY 11973, USA

**Keywords:** rice NIN-like protein 3 (OsNLP3), nitrogen use efficiency (NUE), grain yield, nitrogen availability, rice (*Oryza sativa*)

## Abstract

Nitrogen (N) is an essential macronutrient for crop growth and yield, but excessive application of N fertilizer has caused serious environmental pollution and greatly increased the cost of agricultural production. One of the effective and economical solutions to this dilemma is to improve the N use efficiency (NUE) of crops. Although some components involved in regulating NUE have been identified, the underlying molecular mechanism remains largely elusive in rice. Here we report that the OsNLP3 (NIN-like protein 3) is an important regulator in NUE and grain yield under nitrate-sufficient conditions. Both NUE and grain yield were significantly improved by enhanced *OsNLP3* expression in the field, but reduced in *osnlp3* mutants. The expression of *OsNLP3* responds to both nitrate and ammonium, while OsNLP3 nuclear retention is only induced by nitrate, not by ammonium. OsNLP3 regulates the expression of a set of genes involved in N transport and assimilation by directly binding to the nitrate-responsive *cis*-element in the promoters of these genes. Our study demonstrates that OsNLP3 is significant for the regulation of NUE and grain yield, particularly in nitrate-rich conditions, thus providing a candidate for improving NUE and grain yield in rice.

## Introduction

Nitrogen (N) is an essential macronutrient for plant growth and development and lower N availability in the field limits the crop productivity (Crawford and Forde, 2002; Robertson et al., 2009). N fertilization is one of the key factors improving crop yield and reducing world hunger (Good et al., 2004). To increase crop yield, farmers in many countries excessively apply N fertilizer but less than 50% of the applied fertilizer is utilized by crops, while the leftover is lost to the environment, causing the collateral environmental pollution (Garnett et al., 2009; Mueller et al., 2014; Undurraga et al., 2017). One of the promising solutions to this dilemma is to improve the N use efficiency (NUE) of crops (Hirel et al., 2011; Mandal et al., 2018). NUE, mainly consisting of NUpE (N uptake efficiency) and NUtE (N utilization/assimilation efficiency), is defined by the grain yield per unit of available N in the soil (Xu et al., 2012; Han et al., 2015).

Nitrate is one of the main N sources for plant utilization, and approximately 40% of the total N taken up in paddy grown rice is absorbed as nitrate, due to nitrification in the rhizosphere (Kirk and Kronzucker, 2005). Plants uptake and transport nitrate by four protein families, including low-affinity transporter NPF/NRT1 (peptide transporter family/nitrate transporter1), high-affinity transporter NRT2, CLC (chloride channel), and SLAH (slow anion channel-associated homologues) families (Krapp et al., 2014). Once taken up in the plant, nitrate is reduced to ammonium by the action of NR (nitrate reductase) and NiR (nitrite reductase), then fixed into Glu (glutamate) and Gln (glutamine) by GS (glutamine synthetase) and GOGAT (glutamate synthase) respectively, and thereafter converted to N-containing molecules, such as proteins, nucleotides and hormones (Crawford, 1995; Xu et al., 2012).

Nitrate also act as an important signaling molecule in plants (Camargo et al., 2007; Ho and Tsay, 2010; Undurraga et al., 2017). AtNRT1.1/AtNPF6.3 is a transceptor that functions as a receptor to perceive external nitrate signaling and as a dual-affinity nitrate transporter (Liu et al., 1999; Ho et al., 2009; Hu et al., 2009). After nitrate is sensed by AtNRT1.1, the signal is transduced into the cell through a Ca^2+^-dependent or Ca^2+^-independent pathway (Mu and Luo, 2019). There are also some other proteins, mostly transcription factors (TFs) that were identified participating in the nitrate signaling transduction and regulating N-related metabolism, growth, and development in Arabidopsis, such as AtANR1 (Arabidopsis nitrate regulated 1), AtNRG2 (nitrate regulatory gene 2), AtLBD37/38/39 (lateral organ boundary domain 37/38/39), AtTGA1/4 (TGACG sequence-specific binding protein 1/4), AtAFB3 (auxin signaling F-box 3), and AtbZIP1 (basic leucine zipper 1) (Zhang and Forde, 1998; Rubin et al., 2009; Vidal et al., 2010; Alvarez et al., 2014; Para et al., 2014; Xu et al., 2016). Moreover, several NIN-like proteins (NLPs) act as central transcription factors in nitrogen responses and nitrate signaling (Konishi and Yanagisawa, 2013; Chardin et al., 2014). AtNLP7 acts as a master regulator in nitrate signaling (Konishi and Yanagisawa, 2013; Wang et al., 2018). The knockout mutant *atnlp7* exhibits an impaired PNR (primary nitrate response), which is a rapid nitrate-induced transcriptional response but without protein synthesis (Castaings et al., 2009; Wang et al., 2018). Noteworthily, AtNLP7 orchestrates the early response to nitrate via a special nuclear retention mechanism, which is regulated by nitrate-AtNRT1.1-Ca^2+^-AtCPK10/30/32 (Ca^2+^-dependent protein kinase10/30/32)-AtNLP7 signaling cascade (Marchive et al., 2013; Liu et al., 2017; Alvarez et al., 2020; Oldroyd and Leyser, 2020). In addition, AtNLP7 also plays a central role in plant NUE. The knockout mutant *atnlp7* displays several features of N starvation, whereas overexpression of *AtNLP7* significantly improves the growth and NUE (Castaings et al., 2009; Yu et al., 2016). AtNLP8 acts as a regulator for nitrate-promoted seed germination by reducing ABA content (Yan et al., 2016). Overexpression of the maize *ZmNLP6* and *ZmNLP8* can complement *atnlp7* mutant by restoring nitrate signaling and assimilation (Cao et al., 2017). Furthermore, ZmNLP5 acts as a central hub in a molecular network associated with N metabolism and modulates the nitrogen response in maize (Ge et al., 2020).

Rice is an important food crop for a large part of the world population and serves as a monocot model plant for research. Over the past decade, much progress has been made to enhance NUE in rice. Overexpression of *OsNRT1.1A/OsNPF6.3* improves N utilization and confers high yield and early maturation (Wang et al., 2018). Variation in *OsNRT1.1B/OsNPF6.5* and *OsNR2* (nitrate reductase 2) contributes to nitrate use divergence between rice subspecies (Hu et al., 2015; Gao et al., 2019). Moreover, OsNRT1.1B also acts as a nitrate sensor and interacts with phosphate signaling repressor OsSPX4 (SPX domain containing protein 4) to form a nitrate–OsNRT1.1B– OsSPX4 regulatory module, integrating the nitrogen and phosphorus signaling networks (Hu et al., 2019). OsNAR2.1 (OsNRT2 associate protein) interacts with nitrate transporters OsNRT2.1, OsNRT2.2 and OsNRT2.3a, enable a rapid response to external nitrate concentrations (Yan et al., 2011). OsNPF6.1^HapB^ was trans-activated by the transcription factor OsNAC42 (NAC family transcriptional factor 42), enhancing NUE and grain yield (Tang et al., 2019). Overexpression of *OsPTR9* (peptide transporter 9) improves NUE and grain yield in rice (Fang et al., 2013). Increasing the OsGRF4 (growth-regulating factor 4) abundance in the balance between OsGRF4 and growth inhibitor DELLA improves NUE and grain yield in green revolution varieties (Li et al., 2018). Increased OsNGR5 (nitrogen-mediated tiller growth response 5) activity promotes rice tillering from nitrogen regulation, boosting rice yield and enhancing NUE (Wu et al., 2020).

Although significant progress has been made in understanding nitrate signaling and improving NUE, the regulatory mechanisms remain largely elusive in rice. We recently reported *OsNLP1* and *OsNLP4* that rapidly respond to N availability and improves grain yield and NUE in rice (Alfatih et al., 2020; Wu et al., 2020). Here we report that *OsNLP3*, the closest homolog to Arabidopsis *AtNLP7* in rice, also plays a significant role in nitrate uptake, assimilation and signaling. Similar to *OsNLP1* and *OsNLP4*, *OsNLP3* expression level is significantly induced by N deficiency and repressed by nitrate/ammonium resupply in shoot. However, it does not respond to N starvation but is induced by nitrate/ammonium resupply in the root. OsNLP3 nucleocytosolic shuttling is only responsive to nitrate but not to ammonium. Overexpression of *OsNLP3* significantly improved grain yields and NUE, particularly in high N conditions, whereas *osnlp3* mutants resulted in severe growth reduction, lower grain yield and NUE in field. Our results indicate that *OsNLP3* is a promising candidate to improve NUE and grain yield in rice for sustainable agriculture.

## Results

### OsNLP3 expression responds to N availability

To investigate the response of *OsNLP3* to N availability, we quantified the *OsNLP3* transcript level with different nitrogen treatments in the wild type Zhonghua 11 (ZH11). *OsNLP3* transcript levels in the shoot were gradually induced after shifted to N deficiency (0 N) conditions from normal N supply conditions (NN) and plateaued at 24 hours after the shift. Upon N resupply (either nitrate or ammonium), the transcript levels declined to its pre-induction level in 1 hour while the KCl control treatment did not alter the transcript level (Figure 1A). Interestingly, *OsNLP3* transcript level in the root exhibited a different pattern. In response to N starvation, it slightly increased 1 hour after the shift but gradually declined to its pre-induction level and remained at this low level. Upon N resupply, *OsNLP3* transcript level was induced and peaked 2 hours after the shift, then declined to its pre-induction level (Figure 1B). Our data show that *OsNLP3* is responsive to both nitrate and ammonium while the response patterns are different between shoot and root.

**Figure 1.**
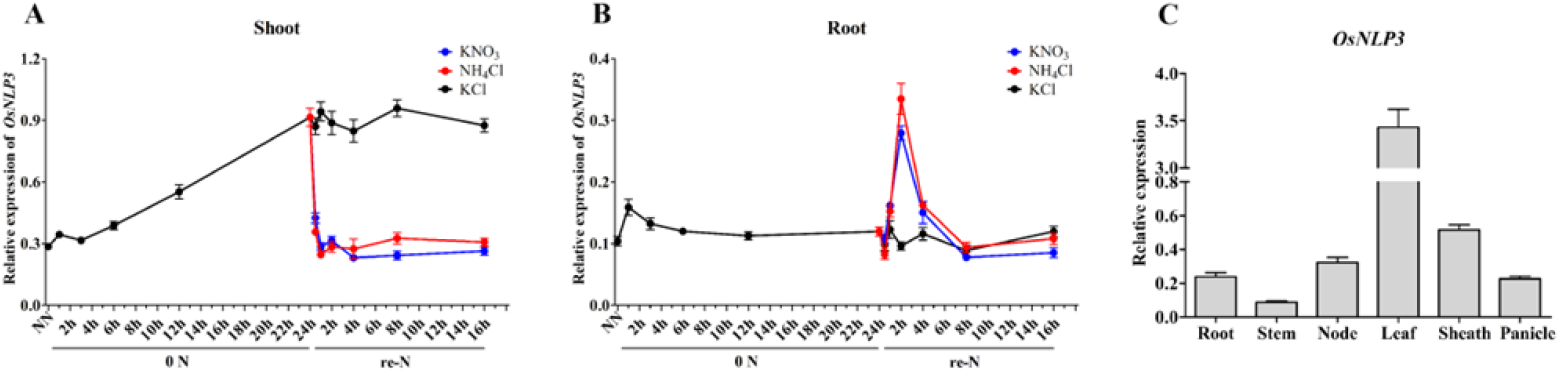
*OsNLP3* expression responds to nitrogen availability. A. Response of *OsNLP3* expression to N availability in shoots of wild type (ZH11). Seedlings were hydroponically grown and treated as described in Materials and Methods (NN, normal nitrogen; 0N, nitrogen starvation; re-N, N resupply). KCl was used as a negative control. Shoots were sampled at indicated time points for RNA analyses with RT-qPCR. Values are the mean ± SD (n = 3). B. Response of *OsNLP3* expression to N availability in root of wild type (ZH11). Roots of the same seedlings as in A were sampled at indicated time points for RNA analyses with RT-qPCR. KCl used as a negative control. Values are the mean ± SD (n = 3). C. Expression of *OsNLP3* in different tissues at flowering stage of wild type (ZH11). RNA was extracted from tissues as indicated and subjected to RT-qPCR analyses. Values are the mean ± SD (n = 3).

RT-qPCR analyses of *OsNLP3* transcript levels in different tissues revealed that it was expressed in all the tissues analyzed but with higher levels in leaves and leaf sheaths (Figure 1C).

### OsNLP3 localization is regulated by nitrate-dependent nuclear retention

To test if the OsNLP3 also have nitrate induced nucleocytosolic shuttling, we generated transgenic rice expressing *pActin1::OsNLP3–GFP* fusion construct in ZH11 background. We found that in seedlings grown in normal N conditions (NN) the subcellular localization of OsNLP3-GFP fusion protein was both in the nucleus and cytosol (Figure 2A), whereas the GFP signals mostly exist in the cytosol in the root of plants grown in N deficiency (0 N) condition (Figure 2B).

**Figure 2.**
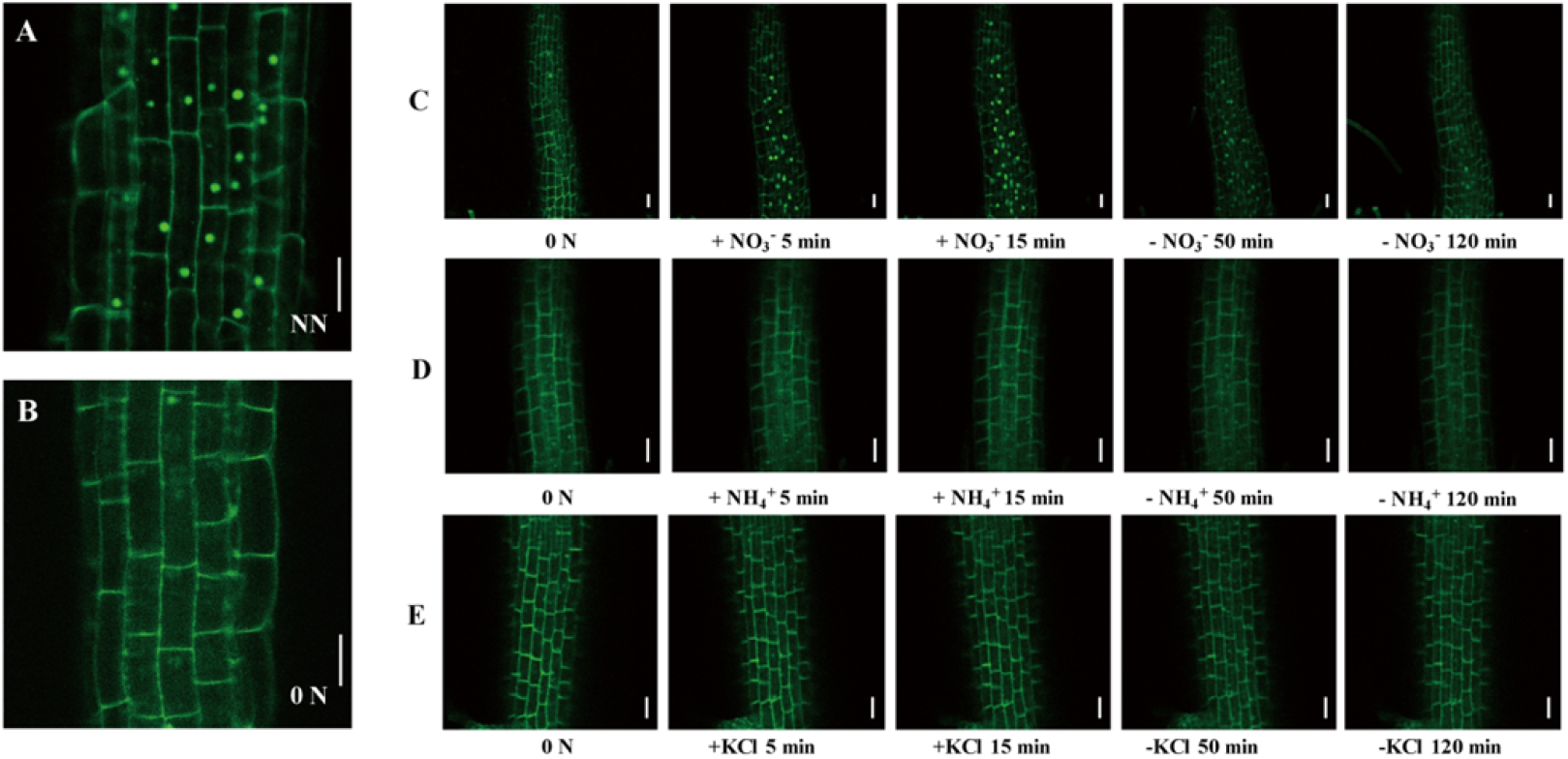
OsNLP3 nuclear retention is only induced by nitrate. A and B. Subcellular localization of OsNLP3-GFP in *pActin1::OsNLP3-GFP* transgenic rice seedlings grown on normal nitrogen medium (A) and N-free medium (B) for 18 days. Bars = 20 μm. C-E. Localization of OsNLP3-GFP in response to nitrate and ammonium. 18-day-old *pActin1::OsNLP3-GFP* seedlings grown on N-free medium (0 N) were transferred to medium with 50 mM KNO_3_ (C) or NH_4_Cl (D) or KCl (E, as a negative control) for 5 min and 15 min, and then transferred to N-free medium for 50 min and 120 min. The images of root cells were captured with a confocal microscope at indicated time points. Bars = 20 μm.

We also examined the nucleocytosolic shuttle of OsNLP3-GFP in response to nitrate and ammonium. In response to resupplied nitrate after N-starvation, OsNLP3–GFP rapidly accumulated in the nucleus within minutes (Figure 2C). This nuclear accumulation was fully reversible when nitrate was withdrawn (Figure 2C). However, when ammonium or KCl was resupplied, the nuclear accumulation of OsNLP3–GFP was not observed (Figure 2D and E), indicating that OsNLP3 nuclear accumulation was specifically responsive to NO_3_^−^ but not to NH_4_^+^ or K^+^.

### OsNLP3 significantly affects seedling growth in nitrate-sufficient conditions

To further investigate the function of OsNLP3 in rice, we generated three loss-of-function mutants (*osnlp3-1, osnlp3-2, osnlp3-3*) with CRISPR/Cas9 system which targeted two different sequences in the first exon of *OsNLP3* (Figure S1A) and two *OsNLP3* overexpression lines (OE-1, OE-10) (Figure S1B). These lines were hydroponically grown in the modified Kimura B solution contained different concentrations of either nitrate or ammonium as the sole N source for 25 days after germination.

Under nitrogen starvation (0 N), seedling growth was obviously retarded for all genotypes without any visible difference (Figure 3A, E-J). Under low N (0.2 mM nitrate as the sole N source) conditions, *osnlp3* mutants only displayed a slight decrease in shoot and root weight, but no obvious difference in shoot length, shoot/root ratio or chlorophyll content of leaves (Figure 3B, E-J), while *OsNLP3* OE plants did not display obvious difference from the wild type except slightly shorter shoot (Figure 3B and I) and higher chlorophyll content (Figure 3J).

**Figure 3.**
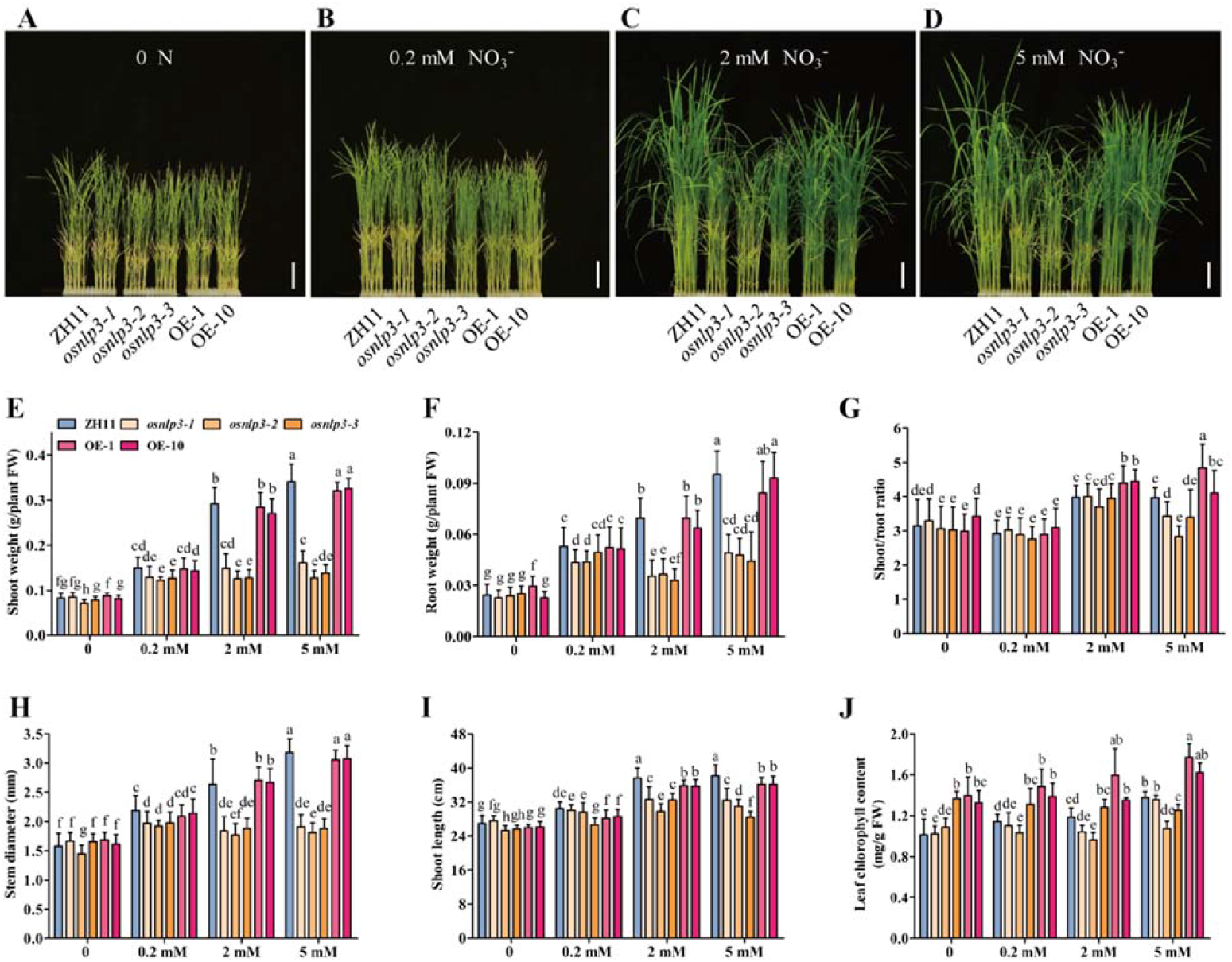
*OsNLP3* affects seedling growth in nitrate-sufficient conditions. A-D. Seedling growth response to nitrate. Wild type (ZH11), mutant (*osnlp3*) and overexpression (OE) plants grown in hydroponic solution with varying amounts of nitrate for 25 days after germination. Bars = 5 cm. E-J. Seedling growth parameters. Shoot fresh weight (E), root fresh weight (F), shoot fresh weight/root fresh weight ratio (G), stem diameter (H), shoot length (I), and leaf chlorophyll content (J) of 25-day-old seedlings were measured. Values are the mean ± SD of 3 independent replications each containing 16 plants per genotype. Different letters denote significant differences evaluated by one-way ANOVA with Tukey’s test (P < 0.05). FW, fresh weight.

By contrast, under normal nitrate (2 mM) and high nitrate (5 mM) conditions the *osnlp3* mutants exhibited constitutive N starvation phenotypes with a biomass (fresh weight) decrease by ~60% in the shoot and ~50% in the root compared to the wild type (Figure 3C-F and H-I). The chlorophyll content of leaves and shoot/root ratio also decreased in *osnlp3* mutants (Figure 3C-D, G and J). The OE lines did not display clear phenotypic difference from the wild type except a slightly higher shoot/root ratio (Figure 3G). Notably, the OE lines were much more green than wild type which support by much higher chlorophyll content under these nitrate conditions (Figure J).

To test the growth response to ammonium, we hydroponically grew different genotypes in culture medium with ammonium as the sole N source. Surprisingly, we did not observe obvious differences among the *osnlp3* mutants, OE lines, and the wild type under low (0.2 mM) or high ammonium (2 mM) conditions (Figure 4A and B, D-I). However, *osnlp3* mutants grown in medium with 1 mM nitrate plus 1 mM ammonium showed a visible retarded growth phenotype compared with wild type (Figure 4C, D-I). These results implicate that OsNLP3 plays a less important role in ammonium utilization.

**Figure 4.**
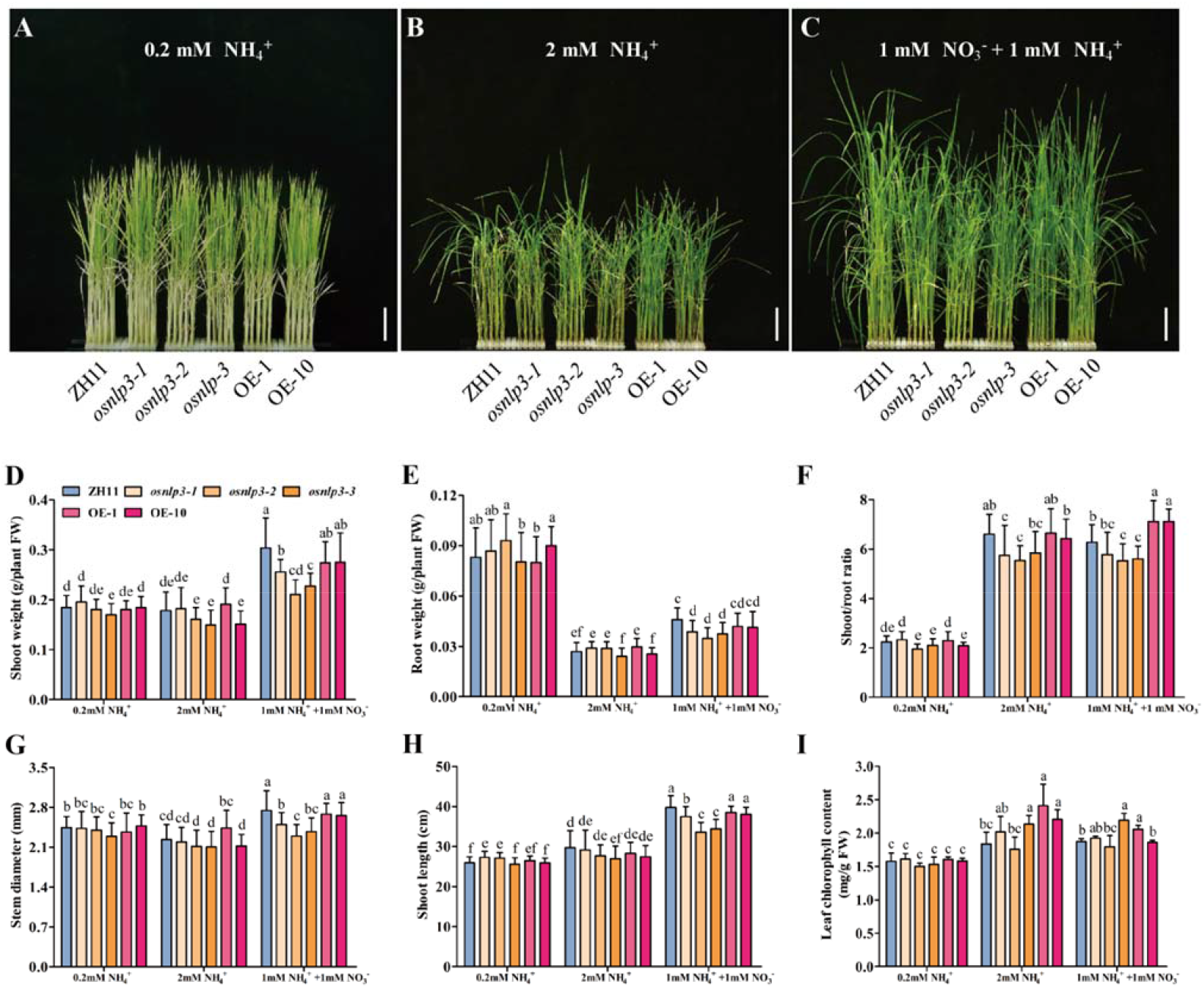
OsNLP3 does not influence seedling growth in response to ammonium. A-C. Seedling growth response to ammonium. Wild type (ZH11), mutant (*osnlp3*) and overexpression (OE) plants grown in hydroponic solution with varying amounts of ammonium for 25 days after germination. Bars = 5 cm. D-I. Seedling growth parameters. Shoot fresh weight (D), root fresh weight (E), shoot fresh weight/root fresh weight ratio (F), stem diameter (G), shoot length (H), and leaf chlorophyll content (I) of 25-day-old seedlings were measured. Values are the mean ± SD of 3 independent replications each containing 16 plants per genotype. Different letters denote significant differences evaluated by one-way ANOVA with Tukey’s test (P < 0.05). FW, fresh weight.

### OsNLP3 positively modulates nitrate uptake and assimilation

The above results of growth response led us to test whether OsNLP3 positively modulates nitrate uptake and assimilation. As expected, the chlorate-sensitivity assay showed that *osnlp3* mutants grew better under hydroponic culture with chlorate, a toxic analog of nitrate, compared with wild type. Inversely, *OsNLP3* OE seedlings exhibited the highest chlorate sensitivity with significantly higher mortality than wild type (Figure 5A and B). Moreover, ^15^N-nitrate or ^15^N-ammonium uptake assays also indicated that nitrate acquisition was reduced in the *osnlp3* mutants while enhanced in *OsNLP3* OE lines, but there was no obvious difference for ammonium absorption (Figure 5C and D). Consistently, the total nitrogen content was decreased in the *osnlp3* mutants and increased in *OsNLP3* OE lines with nitrate as sole nitrogen source, but there was no difference under ammonium conditions (Figure 5E and F). We also measured the free nitrate content and NR activity in the leaves of seedlings hydroponically grown under low (0.2 mM) and normal (2 mM) nitrate conditions. The free nitrate content was higher in *osnlp3* mutants and lower in *OsNLP3* OE lines under normal nitrate, but no obvious difference under low nitrate solution (Figure 5G), indicating that nitrate assimilation is impaired in *osnlp3*, which is supported by significantly reduced NR activity in *osnlp3* mutants and enhanced NR activity in *OsNLP3* OE plants under normal nitrate conditions (Figure 5H). These results indicate that OsNLP3 may directly modulates the uptake and assimilation of nitrate but not ammonium to improve both NUpE and NUtE.

**Figure 5.**
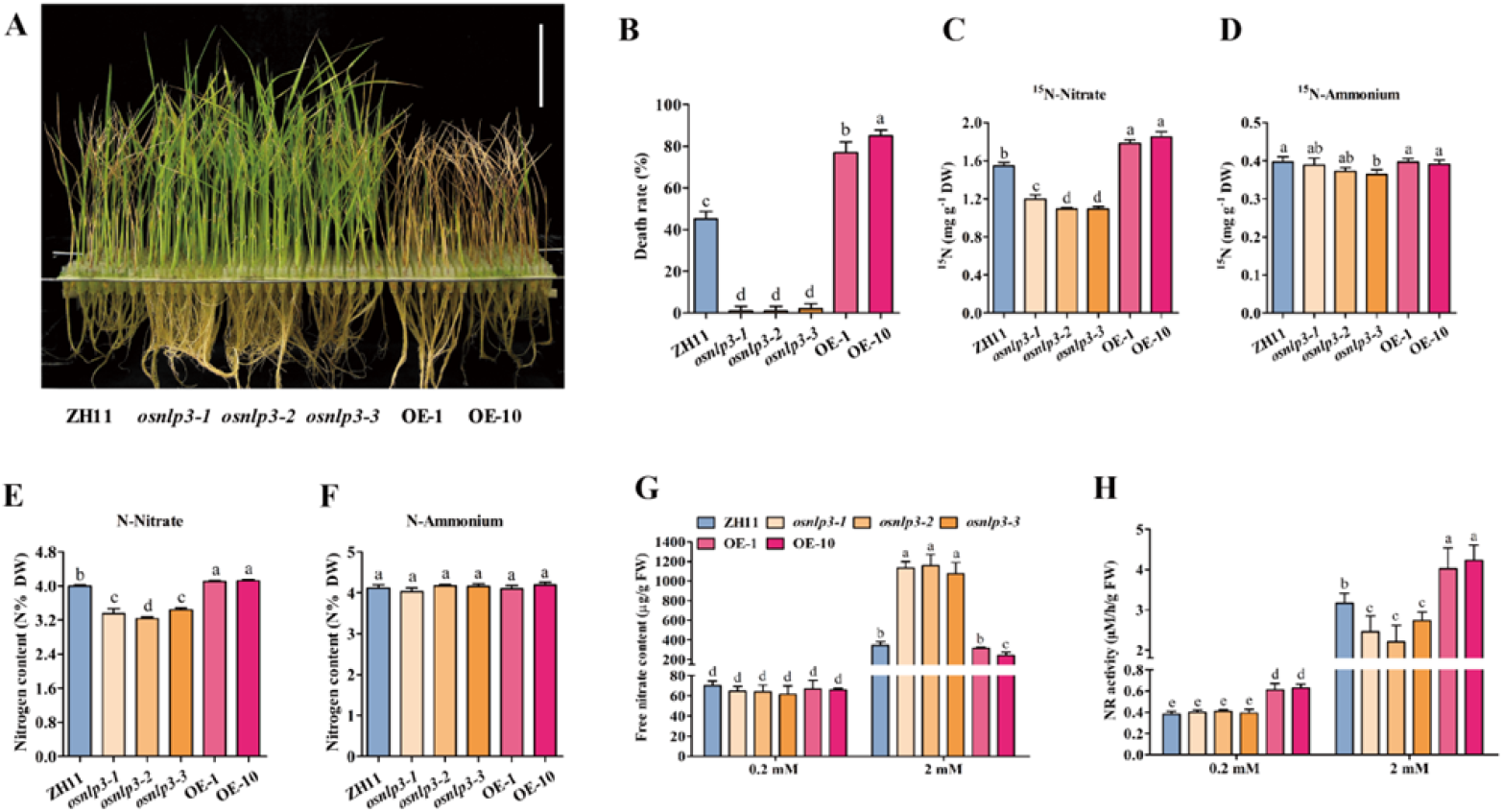
OsNLP3 modulates nitrate uptake and assimilation. A. Chlorate sensitivity assay. Wild type (ZH11), mutant (*osnlp3*) and overexpression (OE) seedlings treated with chlorate as described in Materials and Methods. Bar = 5 cm. B. Chlorate sensitivity. Chlorate sensitivity was evaluated by the death ratio after a chlorate-sensitivity test. Values are the mean ± SD of 3 independent replications each containing 48 plants per genotype. Different letters denote significant differences evaluated by one-way ANOVA with Tukey’s test (P < 0.05). C and D. N uptake assay with ^15^N-nitrate or ^15^N-ammonium. 12-day-old wild type (ZH11), mutant (*osnlp3*) and overexpression (OE) plants were used for N uptake assays as described in Materials and Methods. Values are the mean ± SD of 3 independent replications each containing 6 plants per genotype. Different letters denote significant differences evaluated by one-way ANOVA with Tukey’s test (P < 0.05). DW, dry weight. E and F. Total nitrogen content. 12-day-old wild type (ZH11), mutant (*osnlp3*) and overexpression (OE) seedlings grown in hydroponic solution with 5 mM nitrate or 2 mM ammonium were sampled for total N content analyses as described in Materials and Methods. Different letters denote significant differences evaluated by one-way ANOVA with Tukey’s test (P < 0.05). DW, dry weight. G and H. Free nitrate content and NR activity. Free nitrate content (G) and NR activity (H) in the shoots of 15-day-old wild type (ZH11), mutant (*osnlp3*) and overexpression (OE) plants grown in hydroponic solution with 0. 2 mM and 2 mM nitrate were assayed as described in Materials and Methods. Values are the mean ± SD of 4 independent replications each contained 3 plants per genotype. Different letters denote significant differences evaluated by one-way ANOVA with Tukey’s test (P < 0.05). FW, fresh weight.

### OsNLP3 regulates the expression of nitrate transport, assimilation, and signaling related genes

To further investigate the molecular mechanism of OsNLP3 coordinating nitrate transport, assimilation, and signaling, we measured the expression of genes known to be associated with these processes. Under normal hydroponic cultivation, the genes for primary nitrate assimilation such as *OsNIA1*, *OsNIA2*, *OsNIA3*, and *OsNIR1,* were significantly downregulated in the *osnlp3* mutants and upregulated in *OsNLP3* OE plants (Figure 6A-D). Consistently, the expression of genes related to nitrate uptake and transport such as *OsNRT1.1B*, *OsNRT1.1C*, *OsNRT2.1*, *OsNRT2.3a* and *OsNRT2.4* were down-regulated in the *osnlp3* mutants, and up-regulated in OE plants except *OsNRT1.1C* (Figure 6E-I). Moreover, OsGRF4 was also down regulated in the *osnlp3* mutants and upregulated in OE plants (Figure 6J). These results suggest that OsNLP3, as a transcription factor, may orchestrate the expression of nitrate transport, assimilation, and signaling related genes.

**Figure 6.**
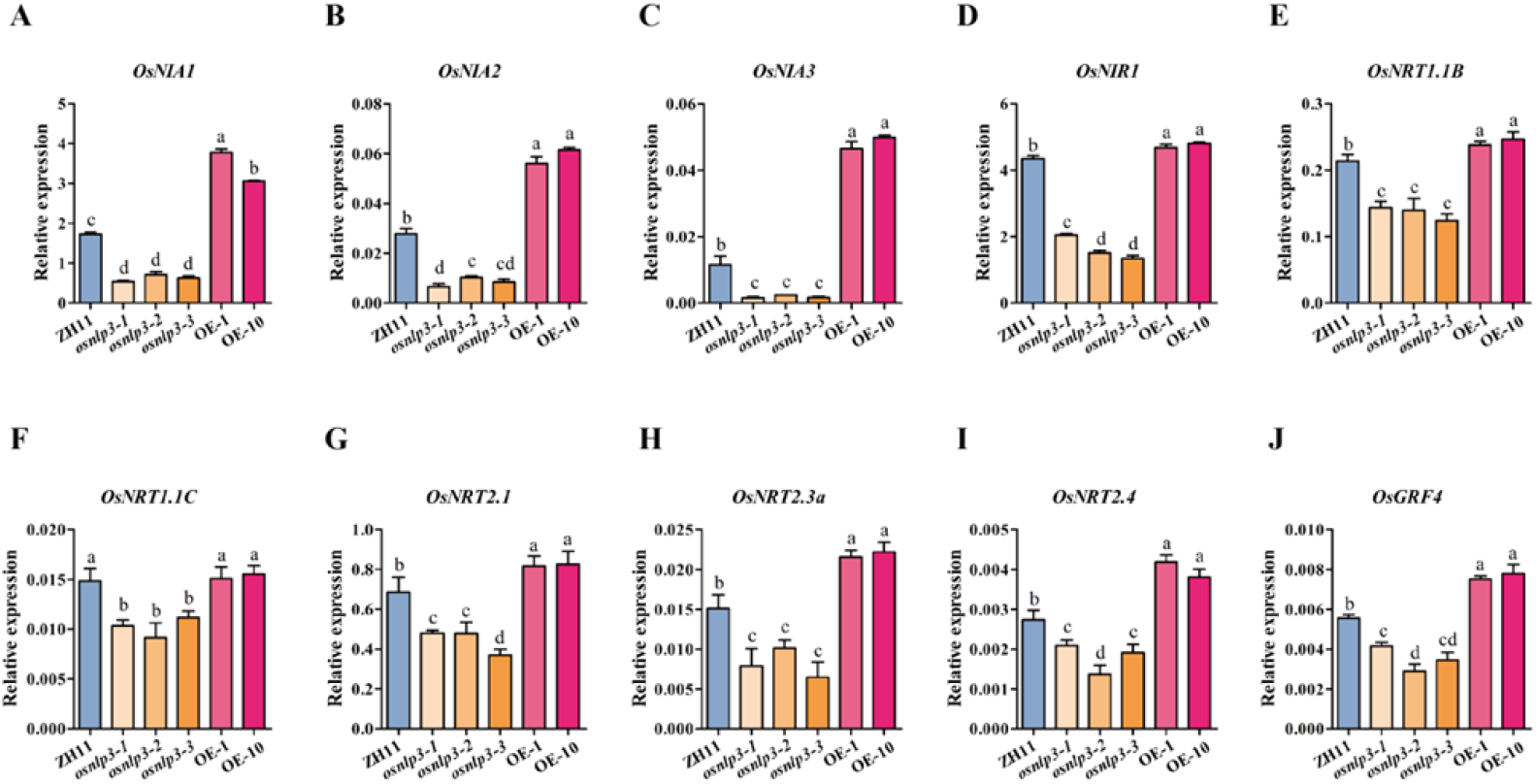
OsNLP3 regulates the expression of nitrate utilization and signaling related genes. A-J. RT-qPCR-based expression analyses of genes involved in nitrate utilization and signaling in wild type (ZH11), mutant (*osnlp3*) and overexpression (OE) plants. 18-day-old plants grown in normal nitrogen medium were used for the analyses. Nitrate reductase genes: *OsNIA1*, *OsNIA2*, and *OsNIA3;* Nitrite reductase gene: *OsNIR1;* Nitrate transporter genes: *OsNRT1.1B, OsNRT1.1C, OsNRT2.1, OsNRT2.3a, OsNRT2.4;* Growth-regulating factor gene: *OsGRF4.* Values are the mean ± SD (*n* = 3). Different letters denote significant differences evaluated by one-way ANOVA with Tukey’s test (P < 0.05).

### OsNLP3 regulates its target genes by directly binding to their promoters

To demonstrate whether OsNLP3 directly modulates the expression of nitrate transport and assimilation related genes, we searched putative NRE-like *cis*-elements in the promoters of the putative target genes regulated by OsNLP3 (Table S1). Those promoters contain one or more NRE-like *cis*-elements. To evaluate whether OsNLP3 directly binds to these NREs, we performed chromatin immunoprecipitation (ChIP) qPCR experiments. ChIP-qPCR results showed that OsNLP3 bound at least one NRE-like *cis-*element in the promoter of N related genes including *OsNIA1*, *OsNIA3*, *OsNRT1.1B*, *OsNRT2.4*, and *OsGRF4* (Figure 7A-E). Yeast-one-hybrid assay (Y1H) further verified these interactions (Figure 7F). These data show that OsNLP3 can directly bind to the promoters of its target genes and potentially regulate their expression.

**Figure 7.**
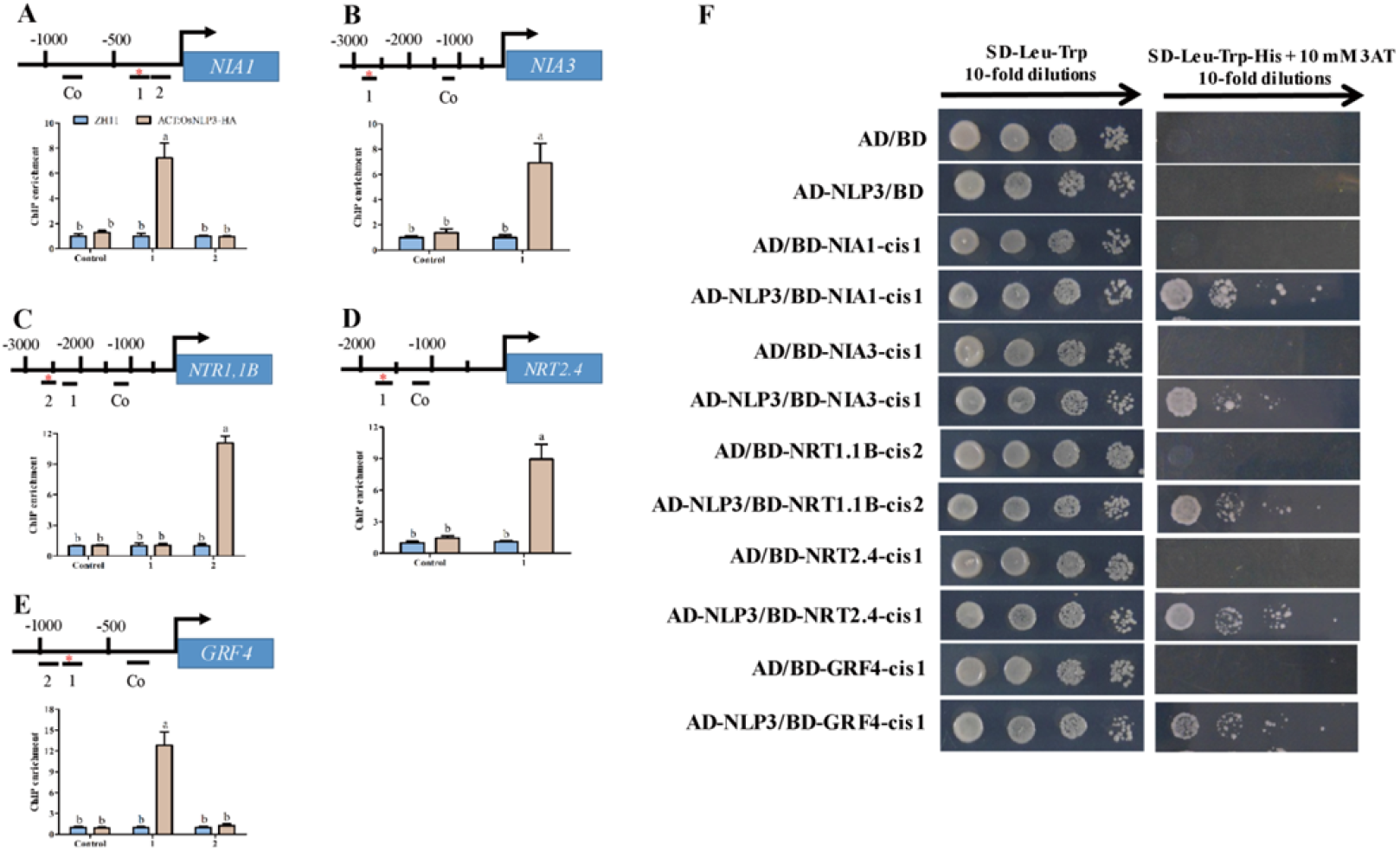
OsNLP3 directly binds to promoters of N-related genes. A-E. ChIP-qPCR assay. HA–OsNLP3 mediated ChIP–qPCR enrichment (relative to wild type) of NRE-like fragments from promoters of *OsNIA1, OsNIA3, OsNRT1.1B, OsNRT2.4, OsGRF4* was performed as described in Materials and methods. Values are the mean ± SD (*n* = 3). Different letters denote significant differences evaluated by one-way ANOVA with Tukey’s test (P < 0.05). F. Yeast-one-hybrid assays. pAD-GAL4-OsNLP3 (AD-NLP3) fusion protein activate the expression of pHIS2 (BD) reporter gene driven by the NRE-like of *OsNIA1, OsNIA3, OsNRT1.1B, OsNRT2.4, OsGRF4*. The empty vector pAD-GAL4 (AD) and pHIS2 (BD) were used as a negative control.

### OsNLP3 overexpression improves NUE and grain yield

To evaluate NUE and grain yield of *osnlp3* mutants and *OsNLP3* OE lines in the field, we designed field trials at two locations (Chengdu, E104°, N30°; Hefei, E117°, N31°). In Chengdu, three nitrogen levels (low nitrogen, LN; normal nitrogen, NN; and high nitrogen, HN) were used to evaluate the performance of each genotype as described in Materials and Methods. Plant height was decreased in *osnlp3* mutants and improved in *OsNLP3* OE plants at all N fertilizer levels compared with wild type (Figure 8A and C).

**Figure 8.**
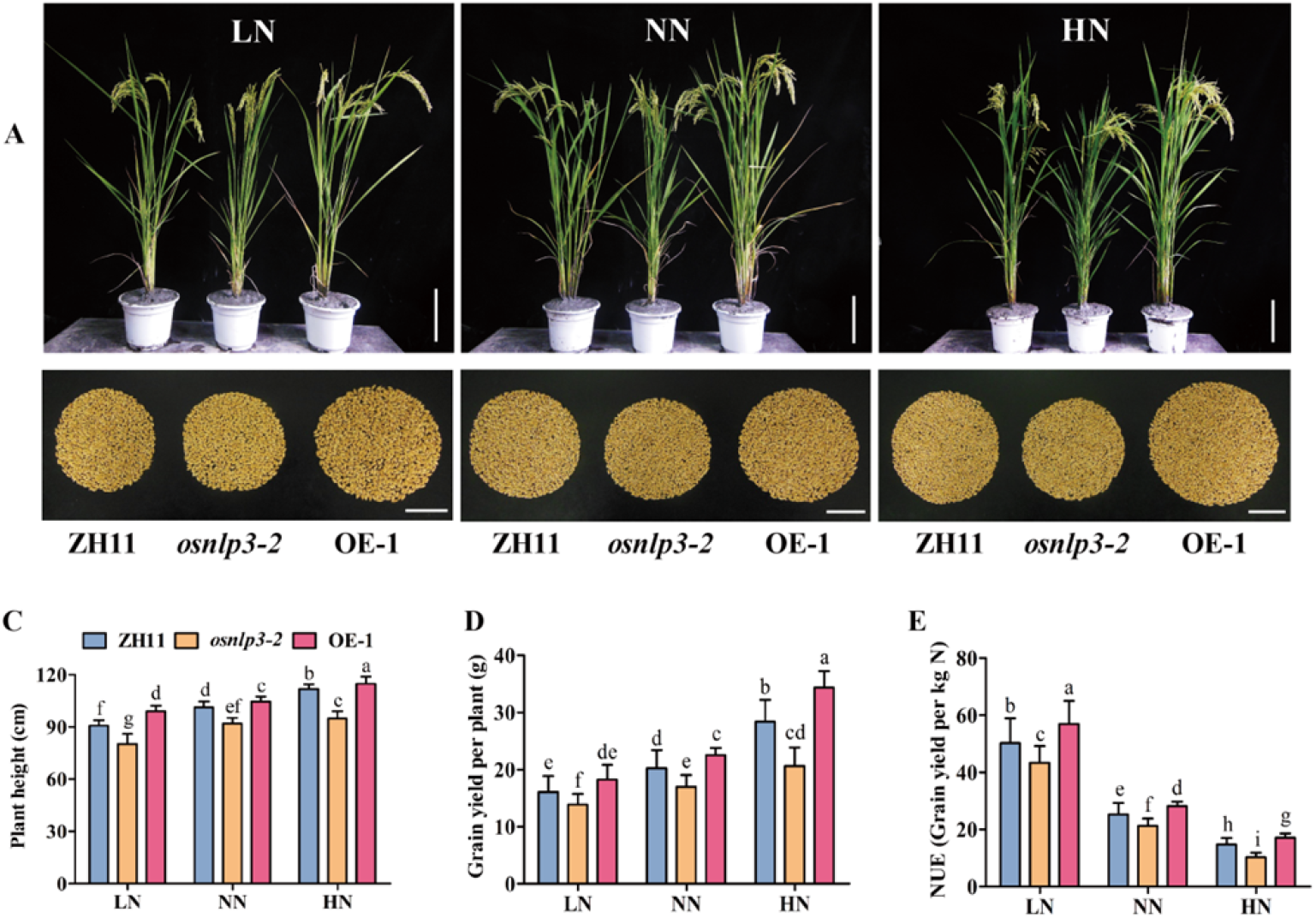
*OsNLP3* overexpression improves NUE and grain yield under different nitrogen levels. A. Morphological phenotypes of wild type (ZH11), mutant (*osnlp3*) and overexpression (OE) grown in the field under low N (LN), normal N (NN) and high N (HN) conditions. A representative plant was dug out from the field and potted for photograph. Bars = 20 cm. B. Total grains per plant of wild type (ZH11), mutant (*osnlp3*) and overexpression (OE) grown in the field under LN, NN and HN conditions. Bars = 5 cm. C-E. Agronomic traits. Plant height (C), grain yield per plant (D) and NUE (E) of wild type (ZH11), mutant (*osnlp3*) and overexpression (OE) plants grown in the field under LN, NN and HN were statistically analyzed. Values are the means ± SD (n=20). Different letters denote significant differences evaluated by one-way ANOVA with Tukey’s test (P < 0.05).

More importantly, *OsNLP3* OE plants improved grain yield per plant and the NUE by ~13% and ~14% under LN, ~11% and ~15% under NN and ~16% and ~20% under HN conditions, respectively. In contrast, the grain yield per plant and the NUE of *osnlp3* mutants exhibited a significant decrease by ~14% and ~15% under LN, ~16% and ~17% under NN and ~30% and ~32% under HN compared to that of the wild type, respectively (Figure 8A-E).

The results of the field tests with normal N level in Hefei were similar to those in Chengdu. Compared with the wild type, plant height was decreased ~8% to 15% in *osnlp3* mutants and increased ~3% to 5% in *OsNLP3* OE plants (Figure 9A and C). Moreover, the grain yield per plant and NUE of *osnlp3* mutants decreased by ~14% - 18% and 19% - 24% while increased by ~13 % - 15% and 16% - 20% in *OsNLP3* OE plants, respectively (Figure 9A-E).

**Figure 9.**
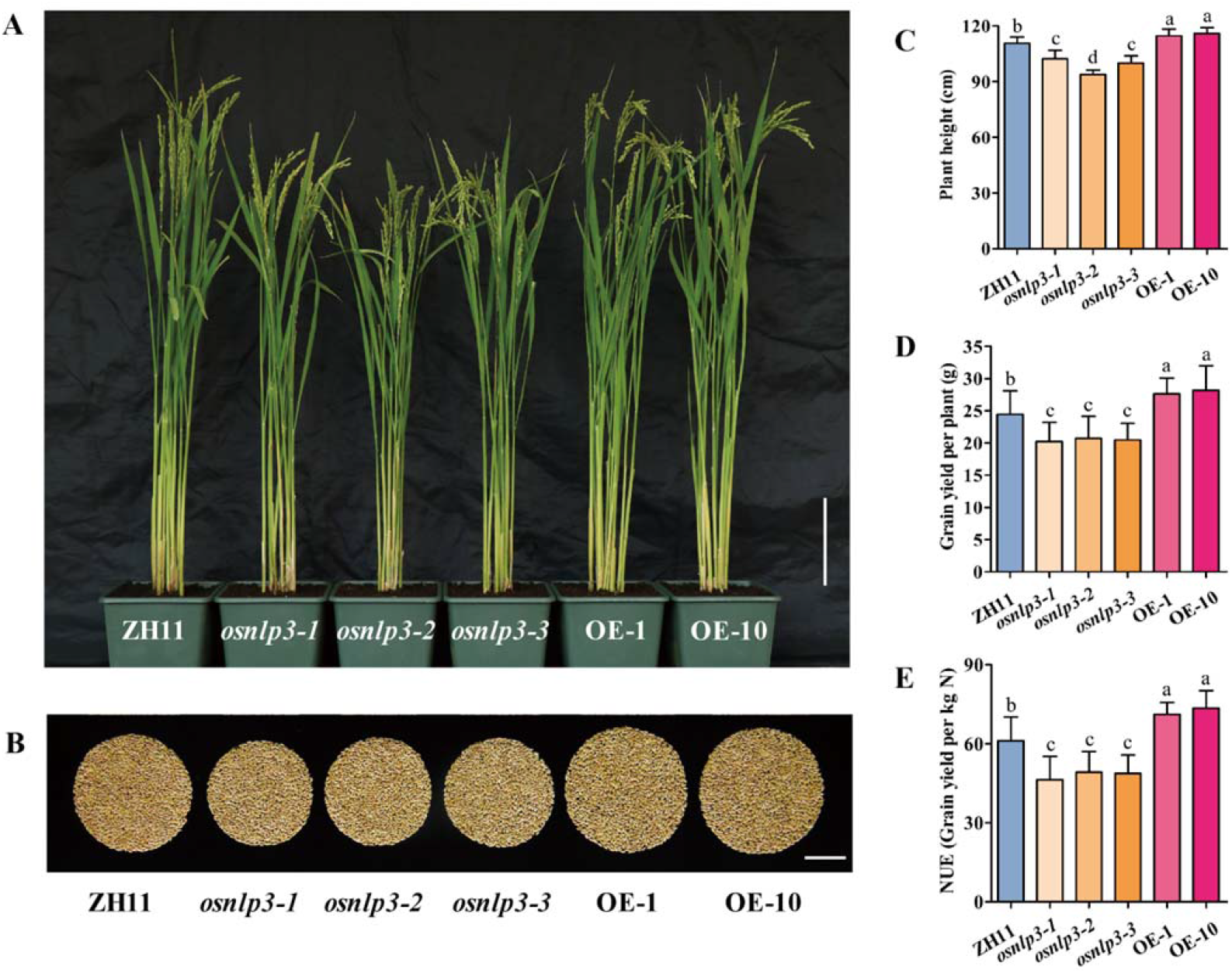
*OsNLP3* overexpression confer to improve NUE and grain yield under normal nitrogen level. A. Morphological phenotypes of wild type (ZH11), mutant (*osnlp3*) and overexpression (OE) grown in the field with normal nitrogen. A representative plant was dug out from the field and potted for photograph. The senescent leaves were removed. Bar = 15 cm. B. Total grains per plant of wild type (ZH11), mutant (*osnlp3*) and overexpression (OE) plants grown in the field with normal nitrogen. Bar = 5 cm. C-E. Agronomic traits. Plant height (C), grain yield per plant (D), and NUE (E) of wild type (ZH11), mutant (*osnlp3*) and overexpression (OE) plants grown in the field with normal nitrogen were statistically analyzed. Values are the means ± SD (n=20). Different letters denote significant differences evaluated by one-way ANOVA with Tukey’s test (P < 0.05).

Taken together, these results clearly demonstrate that OsNLP3 improves NUE and grain yield in the field. In addition, the differences in yield and NUE between different genotypes appears to be less significant under LN conditions but significantly augmented under HN conditions, suggesting that OsNLP3 plays a more important role under HN than LN conditions.

### *OsNLP3* can rescue Arabidopsis *nlp7* and increase biomass

As the most homologous gene to Arabidopsis *NLP7*, we predicted that *OsNLP3* could functionally complement Arabidopsis *nlp7* mutant. The results showed that *OsNLP3* overexpression in Arabidopsis *nlp7* mutant not only rescued the N starvation phenotypes of *nlp7*, but increased biomass compared with wild type under normal (3 mM) and rich (10 mM) nitrate conditions, while no obvious improvement was seen under poor (1 mM) nitrate conditions (Figure S2), consistent with the observations in rice OE lines.

## Discussion

Nitrogen fertilizers are applied excessively in the field to achieve higher yields, especially for green revolution varieties, which is not sustainable and generate serious environmental problems (Han et al., 2015). It has become an urgent issue and a major challenge to reduce fertilizer overuse to protect the environment and increase crop yields to feed the world population. Breeding smart crops that produce acceptable yield with minimum fertilizer input would certainly alleviate the issue. Therefore, improving crop NUE is one of the most economic solutions.

We recently reported *OsNLP1* and *OsNLP4* that improve N utilization and enhance yield and NUE in rice (Alfatih et al., 2020; Wu et al., 2020). In this study, we unveil that OsNLP3 also plays a crucial role in regulating nitrate utilization and NUE in rice. Different from *OsNLP1* and *OsNLP4*, *OsNLP3* shows a different expression pattern in response to N availability in shoot and root. OsNLP3 nuclear localization was specifically and rapidly regulated by nitrate via nuclear retention mechanism but not by ammonium. Moreover, multiple N assimilation genes were significantly down-regulated in *osnlp3* mutants while upregulated in OE lines. More importantly, field tests of the loss-of-function mutants and OE lines clearly demonstrated that OsNLP3 plays a crucial role in NUE and grain yield, particularly in high N conditions.

### Why do shoots and roots show different response pattern to N availability?

OsNLP3 displays an interesting expression pattern with strong induction by N-starvation and repressed by either nitrate or ammonium resupply in the shoot, whereas in the root, it is induced by nitrate or ammonium (Figure 1A-B). This pattern is different from that of *OsNLP1* and *OsNLP4,* which exhibit similar pattern in shoot and root in response to N availability (Alfatih et al., 2020; Wu et al., 2020). It is also different from *AtNLP7* that is not responsive to nitrate (Marchive et al., 2013). The different response pattern to nitrogen in the shoot and root implies that *OsNLP3* may play a more exquisite role to mediate N use and facilitate NUtE in the shoot under N deficiency conditions. In contrast, the induction of *OsNLP3* in the root could rapidly accelerate NUpE when N availability becomes high. This point is supported by higher nitrate absorption (Figure 5) and expression levels of nitrate transport genes (Figure 6) in *OsNLP3* overexpression lines.

### OsNLP3 nucleocytoplasmic shuttle is only responsive to nitrate

OsNLP3 nuclear localization is regulated specifically by nitrate but not by ammonium possibly via the nuclear retention mechanism (Figure 2). This is quite different from OsNLP1, which is localized in the nucleus regardless of the availability of nitrate or ammonium (Alfatih et al., 2020), and different from OsNLP4, whose nuclear localization is responsive to both nitrate and ammonium although much slower and weaker to ammonium (Wu et al., 2020). Considering that AtNLP7 activates the genes involved in PNR via nuclear retention in Arabidopsis (Marchive et al., 2013), we speculate that the transcription factor OsNLP3 maybe also activate PNR and allowing a rapid OsNLP3-dependent genes expression by nuclear retention mechanism in rice. Taken together, given that the nitrate is most readily leached form and spatiotemporal distribution disproportion in field, the swift regulation of OsNLP3 by nitrate via nuclear retention (Figure 2) together with different expression pattern in shoot and root (Figure 1) highlights the crucial role of OsNLP3 in the response to nitrate availability and nitrate signaling pathway.

### OsNLP3 is less important in ammonium utilization

Although *OsNLP3* expression responds to ammonium similarly to nitrate (Figure 1A and B), no obvious phenotypical difference was detected among the wild type, *OsNLP3* OE and mutant lines when grown in medium with ammonium as sole N source (Figure 4). This inconsistence maybe due to the OsNLP3 protein localization in response to ammonium (Figure 2D). Nuclear retention of OsNLP3 only responds to nitrate but not ammonium. OsNLP3-GFP proteins were mainly found in cytosol with hardly any signal in nucleus in plants grown in ammonium medium (Figure 2). Thus, OsNLP3 is unlikely to modulate N assimilation in conditions with ammonium as sole N source. Consistently, chlorate sensitivity assay and ^15^N-ammonium uptake assay revealed that OsNLP3 only affects nitrate uptake rather than ammonium uptake (Figure 5). Similar to *OsNLP3*, *OsNLP1* and *OsNLP4* also respond to both nitrate and ammonium, however, unlike OsNLP3, OsNLP1 proteins constitutively localize in nucleus and OsNLP4 nuclear retention is induced by both nitrate and ammonium (Alfatih et al., 2020; Wu et al., 2020). Consequently, both OsNLP1 and OsNLP4 regulate both nitrate and ammonium assimilation, whereas OsNLP3 mainly functions in nitrate rather than in ammonium utilization.

### OsNLP3 plays more important roles in nitrate-sufficient condition to promote N utilization and improve NUE and yield

Notably, functional characterization of the loss-of-function mutants and OE rice at seedling stage in hydroponic culture demonstrated that OsNLP3 appears to function in promoting nitrate utilization, particularly in sufficient nitrate conditions, but not ammonium (Figure 3, Figure 4). This is supported by chlorate sensitivity assay, ^15^N-uptake assay, total nitrogen content, free nitrate content and NR activity (Figure 5). In contrast, OsNLP1 and OsNLP4 functions in both nitrate and ammonium utilization (Alfatih et al., 2020; Wu et al., 2020). Field test show that grain yield and NUE in mutants decreased ~14% and ~15% under LN, ~30% and ~32% under HN respectively, whereas OE plants increased ~13% and ~14% under LN, ~16% and ~20% under HN, respectively. These results further support that OsNLP3 exhibits stronger functions under N-sufficient conditions than under N-deficient conditions. Inversely, OsNLP1 and OsNLP4 field phenotype is more evident under deficient N conditions than under sufficient N conditions (Alfatih et al., 2020; Wu et al., 2020). The functional divergence between OsNLP3 and OsNLP1/4 in high-N or N-limited conditions suggests that multiple strategies be integrated by OsNLP family members to enhance plant growth under the full range of N statuses.

In conclusion, our study demonstrate that *OsNLP3* plays a crucial role in nitrate utilization and NUE and is a promising candidate to improve NUE and grain yield in rice for sustainable agriculture.

## Materials and Methods

### Plant materials and growth conditions

The rice (*Oryza sativa L.*) *japonica* variety Zhonghua11 (ZH11) was used in this study. Three loss-of-function mutants *osnlp3-1*, *osnlp3-2, osnlp3-3* were obtained through CRISPR-Cas9 technology (Lu et al., 2017) via service provided by Hangzhou Biogle Co., Ltd (Hangzhou, China) and verified by sequencing (Figure S1A). Two *OsNLP3* OE lines (OE-1, OE-10) were generated by inserting the full CDS of *OsNLP3* into the vector of pCB2006 via GATEWAY cloning system (Lei et al., 2007), introduced into ZH11 via *Agrobacterium*-mediated transformation, and verified by RT-PCR analyses (Figure S1B).

For hydroponic culture, seeds were submerged in water at room temperature for 48 h, followed by germination at 37°C. Seeds were then sown in a bottomless 96-well plate that was placed in modified Kimura B solution in a growth chamber with a 14-hour light (30°C)/10-hour dark (28°C) photoperiod, ~300 μmol/m^2^/s light intensity, and ~70% humidity. The modified Kimura B solution contains the following macronutrients (mM): (NH_4_)_2_SO_4_ (0.5), KNO_3_ (1), MgSO_4_·7H_2_O (0.54), CaCl_2_ (0.36), K_2_SO_4_ (0.09), KH_2_PO_4_ (0.18), and Na_2_SiO_3_·9H_2_O (0.7); and micronutrients (μM): MnCl_2_·4H_2_O (9.14), H_3_BO_3_ (46.2), Na_2_MoO_4_ ·2H_2_O (0.56), ZnSO_4_·7H_2_O (0.76), CuSO_4_·5H_2_O (0.32), and Fe(II)-EDTA (40), with the pH adjusted to 5.8.

### Growth response of hydroponically grown seedlings to nitrate and ammonium

For growth response assay of *osnlp3* mutants and *OsNLP3* OE lines (T3 and T4 generation), seedlings were growth in hydroponic solutions with nitrate or ammonium at different concentrations. For 0 N solution, all nitrogen was removed. For 0.2 mM, 2 mM, 5 mM nitrate solutions, all nitrogen was removed and KNO_3_ was added to the indicated concentrations. For 0.2 mM and 2 mM ammonium solutions, all nitrogen was removed and NH_4_Cl was added to the indicated concentrations. For nitrate and ammonium mixed solution, all nitrogen was removed and 1 mM NH_4_Cl plus 1 mM KNO_3_ were added. The nutrient solution for hydroponic culture was renewed every other day. The plants were grown in a growth chamber with a 14-hour light (30°C)/10-hour dark (28°C) photoperiod, ~300 μmol/m^2^/s light intensity, and ~70% humidity for 25 days before sample collection.

### Complementation of Arabidopsis *nlp7-1* with *OsNLP3*

The *Arabidopsis thaliana* ecotype Columbia (Col), *nlp7-1* (SALK_26134C) mutant, and *AtNLP7* OE lines were previously described (Yu et al., 2016). Two *OsNLP3* OE Arabidopsis lines in *nlp7-1* background (*OsNLP3*-OE-4, *OsNLP3*-OE-8) were obtained by *Agrobacterium*-mediated floral-dip method with *p35S::OsNLP3* construct. Plant growth conditions were previously described (Yu et al., 2016). Nitrate medium was modified on Murashige and Skoog (MS) medium with KNO_3_ as sole N source: 10 mM nitrate medium (similar to MS except all nitrogen source was replaced with 9 mM KCl and 10 mM KNO_3_), 3 mM nitrate medium (similar to MS except all nitrogen source was replaced with 16 mM KCl and 3 mM KNO_3_), 1 mM nitrate medium (similar to MS except all nitrogen source was replaced with 18 mM KCl and 1 mM KNO_3_).

### RNA extraction, cDNA preparation, and RT-qPCR

Total RNA was extracted using Trizol reagent (TransGen, Beijing, China, Cat. No. ET101-01) from the indicated tissues of rice plants. One microgram of total RNA was used to synthesize cDNA using Easyscript One-step gDNA Removal and cDNA Synthesis Super Mix reagent kit (TransGen, Beijing, China, Cat. No. AE311-01). qRT-PCR was performed with a StepOne Plus Real Time PCR System by using a SYBR Premix Ex Taq II reagent kit (Vazyme, Nanjing, China, Cat. No. Q311-02). Rice *Actin1* was used as the internal reference. All the primers used are shown in Supplementary Table S2.

### Subcellular localization of OsNLP3-GFP

To investigate the subcellular localization of OsNLP3, the *pActin1::OsNLP3-GFP* fusion constructs was constructed by inserting the full CDS of *OsNLP3* into the pCB2006-*Actin1::GFP* vector via GATEWAY cloning system and transformed into ZH11 (Lei et al., 2007). Transgenic rice seedlings were grown on normal modified Kimura B solution medium or N-free Kimura B solution medium for 18 days.

To observe the nuclear-cytoplasmic shuttling of OsNLP3, we designed nitrate addition and depletion kinetics experiments. Seedlings grown on N-free modified Kimura B solution medium for 18 days, then treated with 50 mM KNO_3_ for 5 min and 15 min, and finally transferred to N-free medium for 50 min and 120 min. Roots were imaged for GFP signals during the time course. For ammonium or KCl (as the control) addition and depletion kinetics experiment, 50 mM NH_4_Cl or 50 mM KCl were used instead of 50 mM KNO_3_ and the rest was the same. Laser scanning confocal imaging of roots was performed using the Zeiss 880 microscope equipped with an argon laser (488nm for green fluorescent protein excitation).

### Response of *OsNLP3* expression to N availability

Rice seedlings of ZH11 were cultured in modified Kimura B solution (NN) for 18 days in growth chamber with a 14-hour light (30°C)/10-hour dark (28°C) photoperiod, ~300 μmol/m^2^/s photon density, and ~70% humidity. For N starvation, seedlings were transferred to N-free modified Kimura B solution (0 N) for 1, 3, 6, 12 and 24 hours, and then transferred to modified Kimura B solution containing 5 mM KNO_3_ or NH_4_Cl (re-N) (5 mM KCl as the control) and grew for 0.5, 1, 2, 4, 8 and 16 hours. Roots and shoots were separately collected at indicated time points for the analyses of *OsNLP3* expression by RT-qPCR.

### Chlorate sensitivity assay

Seedlings were first cultured in modified Kimura B solution containing 2 mM KNO_3_ for 4 days after germination. Seedlings were subsequently treated with 2 mM chlorate for 2 days and allowed to recover in modified Kimura B solution (2 mM KNO_3_) for 5 days. Chlorate sensitivity was calculated as the percent of death. Three replicates were used for chlorate sensitivity assays and each replicate contains 48 seedlings.

### N uptake assay using ^15^N-nitrate or ^15^N-ammonium

^15^N-labeled KNO_3_ (99 atom % ^15^N; Sigma-Aldrich; Cat. No. 335134) or ^15^N-labeled NH_4_Cl (98 atom % ^15^N; Sigma-Aldrich; Cat. No. 299251) were used for N uptake assay. For ^15^N-nitrate uptake assay, rice seedlings were cultured in the modified Kimura B solution with 5 mM KNO_3_ for 12 days. Next, the seedlings were pretreated with the fresh modified Kimura B solution with 5 mM KNO_3_ for 2 hours and then transferred to modified Kimura B solution containing 5 mM ^15^N-KNO_3_ for 24 h. At the end of labeling, the roots were washed for 1 min to remove the ^15^N-KNO_3_ on the root surface. Then the seedlings were dried at 70 °C to a constant weight and ground to powder. For ^15^N-ammonium uptake assays, we followed the same procedure as for nitrate uptake except that 5 mM KNO_3_ and 5 mM ^15^N-KNO_3_ was replaced with 2 mM NH_4_Cl and 2 mM ^15^N-NH_4_Cl, respectively. Finally, the samples were ground and the ^15^N was analyzed using a continuous-flow isotope ratio mass spectrometer (DELTA V Advantage) coupled with an elemental analyzer (EA-HT, Thermo Fisher Scientific, Inc., Bremen, Germany). For each sample, 6 seedlings were collected as a sample, and three biological replicates were used as described (Hu et al., 2015).

### Metabolite analyses and NR activity assay

For chlorophyll content measurement, 25-day-old seedlings grown hydroponically with different N sources and concentrations as phenotyping assays were used. Leaf chlorophyll was extracted by using 80% acetone, and measured by spectrophotometric method as described previously (Yu et al., 2016).

For total N content assay, 12-day-old seedlings cultured hydroponically with 5 mM KNO_3_ or 2mM NH_4_Cl as N sources were collected and dried at 70 °C to a constant weight and ground. Samples were analyzed with an NC analyzer (Vario EL III model, Elementar, Hanau, Germany) according to the manufacturer’s instructions.

For free nitrate content and NR activity assay, 15-day-old seedlings were cultured hydroponically with 0.2 mM /2 mM KNO_3_ as N sources. Free nitrate was extracted in 50 mM HEPES–KOH (pH 7.4), and measured as described previously (Cataldo et al., 1975). The maximum i*n vitro* activity of NR was assayed as previously described (Sylvie et al., 1998).

### Chromatin immunoprecipitation-qPCR

A chromatin immunoprecipitation (ChIP) assay was carried out according to the protocol described previously (Cai et al., 2014). The leaves of one-month-old *pActin1::OsNLP3-HA* transgenic rice plants grown in soil, anti-HA antibodies (Abmart, Shanghai, China), and salmon sperm DNA/protein A agarose beads (Millipore, USA) were used for ChIP experiment. DNA was purified using phenol/ chloroform (1:1, v/v) and precipitated. The enrichments of DNA fragments were quantified by qPCR using specific primers (Table S2). Enriched values were normalized with the level of input DNA.

### Yeast one-hybrid assay

The putative NRE from the flanking sequences of N-relates genes and full CDS of OsNLP3 were cloned into BD vector (pHIS2) and AD vector (pAD-GAL4-2.1), respectively. A yeast one-hybrid assay was conducted according to the procedure described previously (Cai et al., 2014).

### Field evaluation of rice agronomic traits

To investigate the field performance of different *OsNLP3* genotypes, field tests using ZH11, T4 generation of *osnlp3* mutants and *OsNLP3* OE lines were conducted in the paddy field under natural growth conditions during the regular rice growing season in 2019 at two experimental locations: Chengdu (E104°, N30°) and Hefei (E117°, N31°). In Chengdu, three different N concentrations were designed and urea was used as the N source with 80 kg N/hm^2^ for low N, 200 kg N/hm^2^ for normal N and 500 kg N/hm^2^ for high N. The plants were transplanted in 8 rows × 10 plants for each plot and 4 replicate plots were used for each N condition. In Hefei, urea was used as the N source with 100 kg N/hm^2^. The plants were transplanted in 15 rows × 10 plants for each plot and 3 replicate plots were used. For field data collection, the edge lines of each plot were excluded to avoid margin effects.

### Analyses of yield and NUE

All the agronomic traits including plant height, grain yield per plant were collected on a single-plant basis. Detailed methods for measurement of these agronomic traits were described previously (Hu et al., 2015). NUE was defined as the total amount of yield in the form of grain achieved per unit of available N (actual yield per plot / total N application per plot) (Xu et al., 2012).

## Statistical analyses

Statistically significant differences were computed based on the one-way ANOVA.

## Accession Numbers

Sequence data from this article can be found in the Arabidopsis TAIR database (https://www.arabidopsis.org) or Rice Genome Annotation Project (http://rice.plantbiology.msu.edu/) under the following accession numbers: *AtNLP7, AT4G24020; OsNLP3, LOC_Os01g13540; OsNIA1, LOC_Os08g36480; OsNIA2, LOC_Os08g36500; OsNIA3, LOC_Os02g53130; OsNIR1, LOC_Os01g25484; OsNRT1.1B, LOC_Os10g40600; OsNRT1.1C, LOC_Os03g01290; OsNRT2.1, LOC_Os02g02170; OsNRT2.3a, LOC_Os01g50820; OsNRT2.4, LOC_Os01g36720; OsGRF4, LOC_Os02g47280*.

## Acknowledgements

This work was supported by grants from the National Natural Science Foundation of China (grant no. 31572183), the National Key R & D Program of China (grant no. 2016YFD0100701), the Fundamental Research Funds for the Central Universities (grant no. WK6030000122), National Natural Science Foundation of China (grant no. 3157110003), and Ministry of Science and Technology of China (grant no. 2016ZX08005-004-003). Alamin Alfatih is a recipient of CAS-TWAS President’s Fellowship and CAS International Postdoctoral Fellowship.

## Author’s contribution

ZSZ, JW, LHY and CBX designed the experiments. ZSZ. performed experiments and data analysis, and wrote the manuscript. JW, JQX, AA, YJH, YS, LQS, and GYW contributed to performing part of the experiments. SMW, YPW, BHH, GHZ, PQ, SGL contributed to field trials. CBX, JW and LHY revised the manuscript. CBX supervised the project.

## Supporting Information

Figure S1. Verification of the *osnlp3* mutants and *OsNLP3* overexpression lines.

Figure S2. *OsNLP3* overexpression in Arabidopsis *nlp7* increases biomass under nitrate-rich condition.

Table S1. Predicted NRE cis elements in the promoters of N related genes.

Table S2. Primers used in this study.

## Parsed Citations

Alfatih A, Wu J, Zhang ZS, Xia JQ, Jan SU, Yu LH, Xiang CB (2020) Rice NIN-LIKE PROTEIN 1 rapidly responds to nitrogen deficiency and improves yield and nitrogen use efficiency. J Exp Bot 71: 6032–6042 Google Scholar: Author Only Title Only Author and Title

Alvarez JM, Riveras E, Vidal EA, Gras DE, Contreras-Lopez O, Tamayo KP, Aceituno F, Gomez I, Ruffel S, Lejay L, Jordana X, Gutierrez RA (2014) Systems approach identifies TGA1 and TGA4 transcription factors as important regulatory components of the nitrate response of Arabidopsis thaliana roots. Plant J 80: 1–13 Google Scholar: Author Only Title Only Author and Title

Alvarez JM, Schinke AL, Brooks MD, Pasquino A, Leonelli L, Varala K, Safi A, Krouk G, Krapp A, Coruzzi GM (2020) Transient genome-wide interactions of the master transcription factor NLP7 initiate a rapid nitrogen-response cascade. Nat Commun 11: 1157 Google Scholar: Author Only Title Only Author and Title

Cai XT, Xu P, Zhao PX, Liu R, Yu LH, Xiang CB (2014) Arabidopsis ERF109 mediates cross-talk between jasmonic acid and auxin biosynthesis during lateral root formation. Nat Commun 5: 5833 Google Scholar: Author Only Title Only Author and Title

Camargo A, Llamas A, Schnell RA, Higuera JJ, Gonzalez-Ballester D, Lefebvre PA, Fernandez E, Galvan A(2007) Nitrate signaling by the regulatory gene NIT2 in Chlamydomonas. Plant Cell 19: 3491–3503 Google Scholar: Author Only Title Only Author and Title

Cao H, Qi S, Sun M, Li Z, Yang Y, Crawford NM, Wang Y (2017) Overexpression of the Maize ZmNLP6 and ZmNLP8 Can Complement the Arabidopsis Nitrate Regulatory Mutant nlp7 by Restoring Nitrate Signaling and Assimilation. Front Plant Sci 8: 1703 Google Scholar: Author Only Title Only Author and Title

Castaings L, Camargo A, Pocholle D, Gaudon V, Texier Y, Boutet-Mercey S, Taconnat L, Renou JP, Daniel-Vedele F, Fernandez E, Meyer C, Krapp A(2009) The nodule inception-like protein 7 modulates nitrate sensing and metabolism in Arabidopsis. Plant J 57: 426–435 Google Scholar: Author Only Title Only Author and Title

Cataldo DA, Maroon M, Schrader LE, Youngs VL (1975) Rapid colorimetric determination of nitrate in plant tissue by nitration of salicylic acid. Commun Soil Sci Plant Anal 6: 71–80 Google Scholar: Author Only Title Only Author and Title

Chardin C, Girin T, Roudier F, Meyer C, Krapp A(2014) The plant RWP-RK transcription factors: key regulators of nitrogen responses and of gametophyte development. J Exp Bot 65: 5577–5587 Google Scholar: Author Only Title Only Author and Title

Crawford NM (1995) Nitrate: Nutrient and Signal for Plant Growth. Plant Cell 7: 859–868, Google Scholar: Author Only Title Only Author and Title

Crawford NM, Forde BG (2002) Molecular and developmental biology of inorganic nitrogen nutrition. Arabidopsis Book 1: e0011 Google Scholar: Author Only Title Only Author and Title

Fang Z, Xia K, Yang X, Grotemeyer MS, Meier S, Rentsch D, Xu X, Zhang M (2013) Altered expression of the PTR/NRT1 homologue OsPTR9 affects nitrogen utilization efficiency, growth and grain yield in rice. Plant Biotechnol J 11: 446–458 Google Scholar: Author Only Title Only Author and Title

Gao Z, Wang Y, Chen G, Zhang A, Yang S, Shang L, Wang D, Ruan B, Liu C, Jiang H, Dong G, Zhu L, Hu J, Zhang G, Zeng D, Guo L, Xu G, Teng S, Harberd NP, Qian Q (2019) The indica nitrate reductase gene OsNR2 allele enhances rice yield potential and nitrogen use efficiency. Nat Commun 10: 5207 Google Scholar: Author Only Title Only Author and Title

Garnett T, Conn V, Kaiser BN (2009) Root based approaches to improving nitrogen use efficiency in plants. Plant Cell Environ 32: 1272–1283 Google Scholar: Author Only Title Only Author and Title

Ge M, Wang Y, Liu Y, Jiang L, He B, Ning L, Du H, Lv Y, Zhou L, Lin F, Zhang T, Liang S, Lu H, Zhao H (2020) The NIN-like protein 5 (ZmNLP5) transcription factor is involved in modulating the nitrogen response in maize. Plant J 102: 353–368 Google Scholar: Author Only Title Only Author and Title

Good AG, Shrawat AK, Muench DG (2004) Can less yield more? Is reducing nutrient input into the environment compatible with maintaining crop production? Trends Plant Sci 9: 597–605 Google Scholar: Author Only Title Only Author and Title

Han M, Okamoto M, Beatty PH, Rothstein SJ, Good AG (2015) The Genetics of Nitrogen Use Efficiency in Crop Plants. Annu Rev Genet 49: 269–289 Google Scholar: Author Only Title Only Author and Title

Hirel B, Tétu T, Lea PJ, Dubois F (2011) Improving Nitrogen Use Efficiency in Crops for Sustainable Agriculture. Sustainability 3: 1452–1485 Google Scholar: Author Only Title Only Author and Title

Ho CH, Lin SH, Hu HC, Tsay YF (2009) CHL1 functions as a nitrate sensor in plants. Cell 138: 1184–1194 Google Scholar: Author Only Title Only Author and Title

Ho CH, Tsay YF (2010) Nitrate, ammonium, and potassium sensing and signaling. Curr Opin Plant Biol 13: 604–610 Google Scholar: Author Only Title Only Author and Title

Hu B, Jiang Z, Wang W, Qiu Y, Zhang Z, Liu Y, Li A, Gao X, Liu L, Qian Y, Huang X, Yu F, Kang S, Wang Y, Xie J, Cao S, Zhang L, Wang Y, Xie Q, Kopriva S, Chu C (2019) Nitrate-NRT1.1B-SPX4 cascade integrates nitrogen and phosphorus signalling networks in plants. Nat Plants 5: 401–413 Google Scholar: Author Only Title Only Author and Title

Hu B, Wang W, Ou S, Tang J, Li H, Che R, Zhang Z, Chai X, Wang H, Wang Y, Liang C, Liu L, Piao Z, Deng Q, Deng K, Xu C, Liang Y, Zhang L, Li L, Chu C (2015) Variation in NRT1.1B contributes to nitrate-use divergence between rice subspecies. Nat Genet 47: 834–838 Google Scholar: Author Only Title Only Author and Title

Hu HC, Wang YY, Tsay YF (2009) AtCIPK8, a CB L-interacting protein kinase, regulates the low-affinity phase of the primary nitrate response. Plant J 57: 264–278 Google Scholar: Author Only Title Only Author and Title

Kirk GJ, Kronzucker HJ (2005) The potential for nitrification and nitrate uptake in the rhizosphere of wetland plants: a modelling study. Annals Botany 96: 639–646 Google Scholar: Author Only Title Only Author and Title

Konishi M, Yanagisawa S (2013) Arabidopsis NIN-like transcription factors have a central role in nitrate signalling. Nat Commun 4: 1617 Google Scholar: Author Only Title Only Author and Title

Krapp A, David LC, Chardin C, Girin T, Marmagne A, Leprince AS, Chaillou S, Ferrario-Mery S, Meyer C, Daniel-Vedele F (2014) Nitrate transport and signalling in Arabidopsis. J Exp Bot 65: 789–798 Google Scholar: Author Only Title Only Author and Title

Lei Z-Y, Zhao P, Cao M-J, Cui R, Chen X, Xiong L-Z, Zhang Q-F, Oliver DJ, Xiang C-B (2007) High-throughput Binary Vectors for Plant Gene Function Analysis. J Integr Plant Biol 49 (4): 556−567 Google Scholar: Author Only Title Only Author and Title

Li S, Tian Y, Wu K, Ye Y, Yu J, Zhang J, Liu Q, Hu M, Li H, Tong Y, Harberd NP, Fu X (2018) Modulating plant growth-metabolism coordination for sustainable agriculture. Nature 560: 595–600 Google Scholar: Author Only Title Only Author and Title

Liu K-H, Huang C-Y, Tsay Y-F (1999) CHL1 is a dual-affinity nitrate transporter of Arabidopsis involved in multiple phases of nitrate uptake. Plant Cell 11: 865–874 Google Scholar: Author Only Title Only Author and Title

Liu KH, Niu Y, Konishi M, Wu Y, Du H, Sun Chung H, Li L, Boudsocq M, McCormack M, Maekawa S, Ishida T, Zhang C, Shokat K, Yanagisawa S, Sheen J (2017) Discovery of nitrate-CPK-NLP signalling in central nutrient-growth networks. Nature 545: 311–316 Google Scholar: Author Only Title Only Author and Title

Lu Y, Ye X, Guo R, Huang J, Wang W, Tang J, Tan L, Zhu JK, Chu C, Qian Y (2017) Genome-wide Targeted Mutagenesis in Rice Using the CRISPR/Cas9 System. Mol Plant 10: 1242–1245 Google Scholar: Author Only Title Only Author and Title

Mandal VK, Sharma N, Raghuram N (2018) Molecular Targets for Improvement of Crop Nitrogen Use Efficiency: Current and Emerging Options. In A Shrawat, A Zayed, DA Lightfoot, eds, Engineering Nitrogen Utilization in Crop Plants. Springer New York, pp 77–93 Google Scholar: Author Only Title Only Author and Title

Marchive C, Roudier F, Castaings L, Brehaut V, Blondet E, Colot V, Meyer C, Krapp A(2013) Nuclear retention of the transcription factor NLP7 orchestrates the early response to nitrate in plants. Nat Commun 4: 1713 Google Scholar: Author Only Title Only Author and Title

Mu X, Luo J (2019) Evolutionary analyses of NIN-like proteins in plants and their roles in nitrate signaling. Cellular Mol Life Sci 76: 3753–3764 Google Scholar: Author Only Title Only Author and Title

Mueller ND, West PC, Gerber JS, MacDonald GK, Polasky S, Foley JA(2014) Atradeoff frontier for global nitrogen use and cereal production. Environ Res Lett 9: 054002 Google Scholar: Author Only Title Only Author and Title

Oldroyd GED, Leyser O (2020) Aplant’s diet, surviving in a variable nutrient environment. Science 368 Google Scholar: Author Only Title Only Author and Title

Para A, Li Y, Marshall-Colon A, Varala K, Francoeur NJ, Moran TM, Edwards MB, Hackley C, Bargmann BO, Birnbaum KD, McCombie WR, Krouk G, Coruzzi GM (2014) Hit-and-run transcriptional control by bZIP1 mediates rapid nutrient signaling in Arabidopsis. Proc Natl Acad Sci USA 111: 10371–10376 Google Scholar: Author Only Title Only Author and Title

Robertson GP, Vitousek PM, resources (2009) Nitrogen in agriculture: balancing the cost of an essential resource. Annu Rev Environ Resour 34: 97–125 Google Scholar: Author Only Title Only Author and Title

Rubin G, Tohge T, Matsuda F, Saito K, Scheible WR (2009) Members of the LBD family of transcription factors repress anthocyanin synthesis and affect additional nitrogen responses in Arabidopsis. Plant Cell 21: 3567–3584 Google Scholar: Author Only Title Only Author and Title

Sylvie FM, Valadier MH, Foyer CH (1998) Overexpression of nitrate reductase in tobacco delays drought-induced decreases in nitrate reductase activity and mRNA. Plant Physiol 117: 293–302 Google Scholar: Author Only Title Only Author and Title

Tang W, Ye J, Yao X, Zhao P, Xuan W, Tian Y, Zhang Y, Xu S, An H, Chen G, Yu J, Wu W, Ge Y, Liu X, Li J, Zhang H, Zhao Y, Yang B, Jiang X, Peng C, Zhou C, Terzaghi W, Wang C, Wan J (2019) Genome-wide associated study identifies NAC42-activated nitrate transporter conferring high nitrogen use efficiency in rice. Nat Commun 10: 5279 Google Scholar: Author Only Title Only Author and Title

Undurraga SF, Ibarra-Henríquez C, Fredes I, Álvarez JM, Gutiérrez R (2017) Nitrate signaling and early responses in Arabidopsis roots. J Exp Bot 68: 2541–2551 Google Scholar: Author Only Title Only Author and Title

Vidal EA, Araus V, Lu C, Parry G, Green PJ, Coruzzi GM, Gutierrez RA(2010) Nitrate-responsive miR393/AFB3 regulatory module controls root system architecture in Arabidopsis thaliana. Proc Natl Acad Sci USA 107: 4477–4482 Google Scholar: Author Only Title Only Author and Title

Wang W, Hu B, Yuan D, Liu Y, Che R, Hu Y, Ou S, Liu Y, Zhang Z, Wang H, Li H, Jiang Z, Zhang Z, Gao X, Qiu Y, Meng X, Liu Y, Bai Y, Liang Y, Wang Y, Zhang L, Li L, Sodmergen, Jing H, Li J, Chu C (2018) Expression of the Nitrate Transporter Gene OsNRT1.1A/OsNPF6.3 Confers High Yield and Early Maturation in Rice. Plant Cell 30: 638–651 Google Scholar: Author Only Title Only Author and Title

Wang YY, Cheng YH, Chen KE, Tsay YF (2018) Nitrate Transport, Signaling, and Use Efficiency. Annu Rev Plant Biol 69: 85–122 Google Scholar: Author Only Title Only Author and Title

Wu J, Zhang ZS, Xia JQ, Alfatih A, Song Y, Huang YJ, Wan GY, Sun LQ, Tang H, Liu Y, Wang SM, Zhu QS, Qin P, Wang YP, Li SG, Mao CZ, Zhang GQ, Chu C, Yu LH, Xiang CB (2020) Rice NIN-LIKE PROTEIN 4 plays a pivotal role in nitrogen use efficiency. Plant Biotechnol J Google Scholar: Author Only Title Only Author and Title

Wu K, Wang S, Song W, Zhang J, Wang Y, Liu Q, Yu J, Ye Y, Li S, Chen J, Zhao Y, Wang J, Wu X, Wang M, Zhang Y, Liu B, Wu Y, Harberd NP, Fu X (2020) Enhanced sustainable green revolution yield via nitrogen-responsive chromatin modulation in rice. Science 367 Google Scholar: Author Only Title Only Author and Title

Xu G, Fan X, Miller AJ (2012) Plant nitrogen assimilation and use efficiency. Annu Rev Plant Biol 63: 153–182 Google Scholar: Author Only Title Only Author and Title

Xu N, Wang R, Zhao L, Zhang C, Li Z, Lei Z, Liu F, Guan P, Chu Z, Crawford NM, Wang Y (2016) The Arabidopsis NRG2 Protein Mediates Nitrate Signaling and Interacts with and Regulates Key Nitrate Regulators. Plant Cell 28: 485–504 Google Scholar: Author Only Title Only Author and Title

Yan D, Easwaran V, Chau V, Okamoto M, Ierullo M, Kimura M, Endo A, Yano R, Pasha A, Gong Y, Bi YM, Provart N, Guttman D, Krapp A, Rothstein SJ, Nambara E (2016) NIN-like protein 8 is a master regulator of nitrate-promoted seed germination in Arabidopsis. Nat Commun 7: 13179 Google Scholar: Author Only Title Only Author and Title

Yan M, Fan X, Feng H, Miller AJ, Shen Q, Xu G (2011) Rice OsNAR2.1 interacts with OsNRT2.1, OsNRT2.2 and OsNRT2.3a nitrate transporters to provide uptake over high and low concentration ranges. Plant Cell Environ 34: 1360–1372 Google Scholar: Author Only Title Only Author and Title

Yu LH, Wu J, Tang H, Yuan Y, Wang SM, Wang YP, Zhu QS, Li SG, Xiang CB (2016) Overexpression of Arabidopsis NLP7 improves plant growth under both nitrogen-limiting and -sufficient conditions by enhancing nitrogen and carbon assimilation. Scientific Rep 6: 27795 Google Scholar: Author Only Title Only Author and Title

Zhang H, Forde BG (1998) An Arabidopsis MADS box gene that controls nutrient-induced changes in root architecture. Science 279: 407–409 Google Scholar: Author Only Title Only Author and Title

